# Continuous addition of progenitors forms the cardiac ventricle in zebrafish

**DOI:** 10.1101/230649

**Authors:** Anastasia Felker, Karin D. Prummel, Anne M. Merks, Michaela Mickoleit, Eline C. Brombacher, Jan Huisken, Daniela Panáková, Christian Mosimann

## Abstract

The vertebrate heart develops from several progenitor lineages. After early-differentiating first heart field (FHF) progenitors form the linear heart tube, late-differentiating second heart field (SHF) progenitors extend atrium, ventricle, and form the inflow and outflow tracts (IFT/OFT). However, the position and migration of late-differentiating progenitors during heart formation remains unclear. Here, we tracked zebrafish heart development using transgenics based on the cardiopharyngeal transcription factor gene *tbx1*. Live-imaging uncovered a *tbx1* reporter-expressing cell sheath that from anterior lateral plate mesoderm continuously disseminates towards the forming heart tube. High-speed imaging and optogenetic lineage tracing corroborated that the zebrafish ventricle forms through continuous addition from the undifferentiated progenitor sheath followed by late-phase accrual of the bulbus arteriosus (BA). FGF inhibition during sheath migration reduced ventricle size and abolished BA formation, refining the window of FGF action during OFT formation. Our findings consolidate previous end-point analyses and establish zebrafish ventricle formation as a continuous process.

## Introduction

Vertebrate cardiomyocytes derive from an early versus a late differentiating progenitor pool that can be divided within the anterior lateral plate mesoderm (ALPM) into the FHF and SHF (Buckingham et al., 2005). After the early-differentiating FHF assembles the linear heart tube that in mammals forms the left ventricle and parts of the atria, the late-differentiating SHF contributes to the atria, right ventricle, and OFT (Kelly, 2012; Kelly et al., 2001; Mjaatvedt et al., 2001; Tzahor and Evans, 2011). In mice, the SHF forms within medial and posterior epithelial-like field in the splanchnic ALPM on either side of the FHF-derived cardiac crescent and is detectable through expression of markers including *Fgf10* and *Isl1* (Cai et al., 2003; Francou et al., 2017; Kelly et al., 2001). Developmental defects in SHF contribution to the heart lead to a broad variety of congenital heart defects affecting the arterial pole(Srivastava and Olson, 2000). The original purpose of a second late-forming wave of myocardium remains unknown. An emerging concept places the SHF in close developmental and evolutionary lineage relationship with pharyngeal and head muscle progenitors: in this context, the cardiopharyngeal field (CPF) is defined across chordate species as a domain within the splanchnic or pharyngeal ALPM harboring SHF and branchiomeric progenitor cells (Diogo et al., 2015). Consequently, a key aspect of SHF is its cardiac specification coordinated with head muscle differentiation (Wang et al., 2017).

The basic mechanisms of vertebrate heart formation are evolutionarily conserved. Typical for teleosts, the zebrafish heart consists of two chambers, an atrium and a ventricle (Figure 1A). At the arterial pole, the myocardium transitions into laminin-rich myocardium referred to as conus arteriosus (CA) followed by the elastic bulbus arteriosus (BA) that functions as smooth muscle-based pressure capacitator similar to the mammalian aortic arch (Grimes and Kirby, 2009; Moriyama et al., 2016); the CA together with the BA are commonly, but not consistently, defined as OFT (Grimes and Kirby, 2009; Grimes et al., 2006) (Figure 1A). Cardiac progenitors become detectable in the zebrafish ALPM by bilateral expression of several conserved cardiac transcription factors, including *nkx2.5, hand2*, and *gata4/5* (Chen and Fishman, 1996; Reiter et al., 1999; Schoenebeck et al., 2007; Yelon et al., 2000). By 14-18 hours post-fertilization (hpf), bilateral progenitors condense at the midline as cardiac disc that forms the cardiac cone; the subsequently emerging linear heart tube consists of endocardium and surrounding early differentiating cardiomyocytes referred to as FHF myocardium, discernible at 16-18 hpf by differentiation markers including *myl7* (Bakkers, 2011; Fishman and Chien, 1997; Stainier et al., 1993). Myocardial expression of *drl*-based transgenes selectively marks FHF-derived cardiomyocytes from late somitogenesis (Mosimann et al., 2015). Starting from 26 hpf, akin to the mammalian heart, a late-differentiating wave of prospective cardiomyocytes and smooth muscle cells extends the venous IFT and arterial OFT of the beating zebrafish heart, referred to as SHF lineage (Hami et al., 2011; Lazic and Scott, 2011; Zhou et al., 2011) (Figure 1A). The zebrafish heart therefore recapitulates key processes of multi-chambered heart formation (Bakkers, 2011).

**Figure 1:**
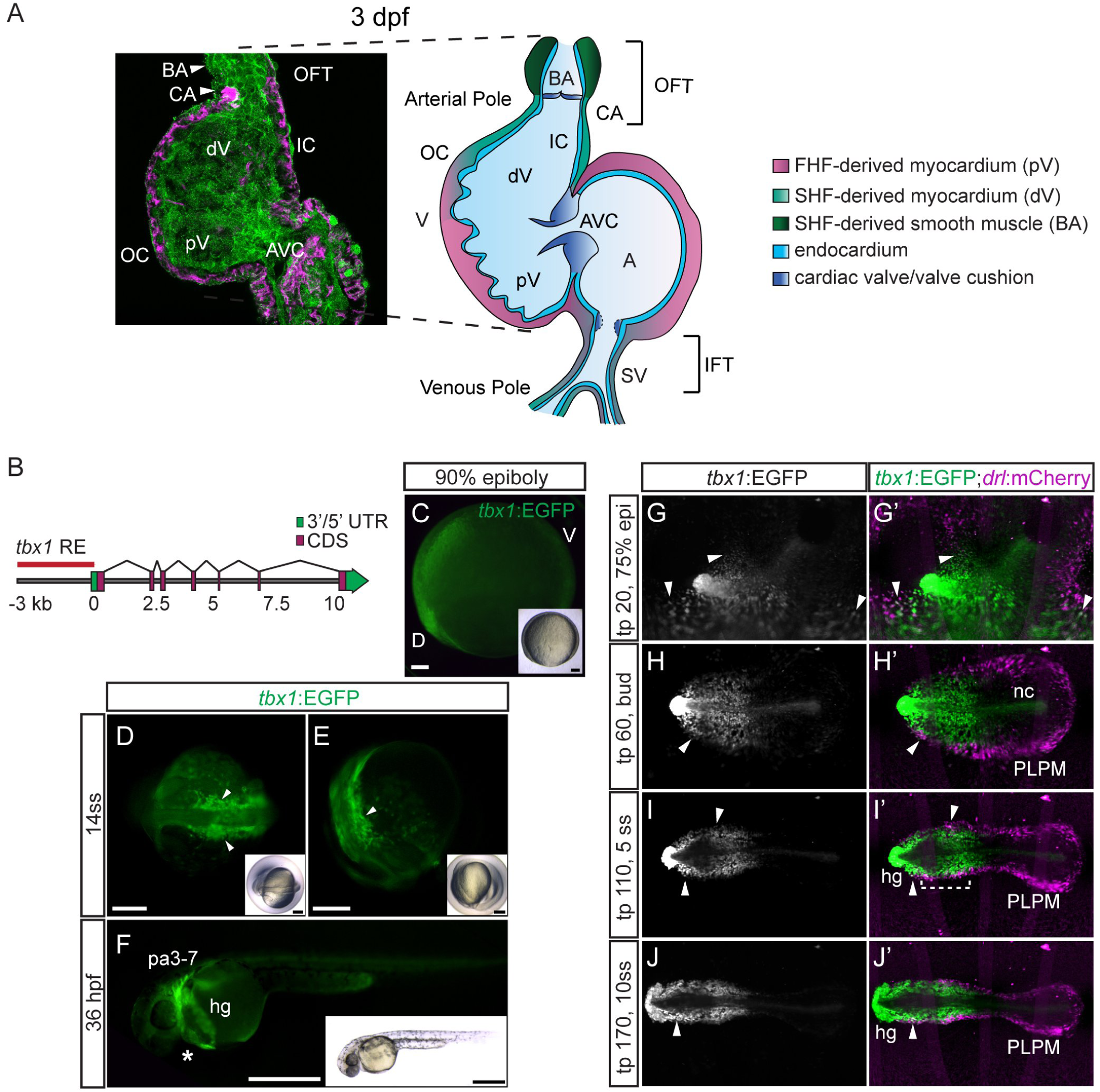
*tbx1* reporter expression and lineage contribution in the cardiopharyngeal and ALPM field. (**A**) Confocal Z-projection and schematic representation of a 72 hpf zebrafish heart with two chambers, the ventricle (V) and atrium (A) separated by a valve at the atrioventricular canal (AVC). The isolated heart is stained for MHC marking the myocardium (magenta) and a-PKC marking all cells (green). The FHF-assigned myocardium contains the proximal ventricle (pV) and the majority of the atrium (A), SHF-assigned myocardium forms the distal ventricle (dV) and outflow tract (OFT), magenta and green in the schematic, respectively. The OFT includes the conus arteriosus (CA), comprising the myocardial connection of the ventricle to the bulbus arteriosus (BA), and the smooth muscular BA itself. The lineage contributions to the sinus venosus (SV)/inflow tract (IFT) and developmental timing of IFT valve formation remain unresolved. IC, inner curvature; OC, outer curvature. (**B**) Genomic locus of the zebrafish *tbx1* gene; red arrow indicates 3.2 kb cis-regulatory region amplified to drive reporter transgenics. (**C**) tbx1:EGFP reporter transgene expression in a dorsal/anterior field during gastrulation (90% epiboly, dorsal/anterior to the left). (**D,E**) *tbx1*:EGFP at 14 ss; dorsal (**D**) and lateral views (**E**) of the prospective cardiopharyngeal field (CPF, arrowheads). (**F**) Lateral view of a 36 hpf *tbx1:EGFP* embryo with EGFP expression in the CPF-derived pharyngeal arches (pa3-7) and heart (asterisk). EGFP expression also marks the prechordal mesoderm-derived hatching gland (hg), insets depict bright-field. (**G-J**) Mercator projection of representative stages from panoramic SPIM-imaged *tbx1:EGFP;drl:mCherry* double-positive transgenic embryos; dorsal views. *tbx1*:EGFP expression is confined to the anterior of the embryo, with no EGFP signal in the posterior LPM (PLPM). Expression in the notochord (nc) and hatching gland (hg) are likely related to early prechordal plate activity of the reporter. Note double-positive cells at the outermost domain of the *tbx1*:EGFP-positive anterior cell population (arrowheads and bracket). Scale bars 100 μm (**C**), 200 μm (**D,E**), 500 μm (**F**). (**D-J**) anterior to the left.

Lineage tracing, several molecular markers including *ltbp3, isl1*, and *mef2c*, or the maturation speed of fluorescent reporters have been used to describe late differentiating SHF-like myocardium (Hami et al., 2011; Lazic and Scott, 2011; de Pater et al., 2009; Witzel et al., 2017, 2012; Zhou et al., 2011). Nonetheless, the lineage separation and developmental connection between the ventricular myocardium and OFT structures remains vaguely defined. Further, the position of SHF-assigned cells during formation of the linear heart tube has remained uncertain. Genetic lineage tracing has shown that both distal ventricle and OFT derive from *nkx2.5*- and *ltbp3*-reporter-expressing cells (Guner-Ataman et al., 2013; Zhou et al., 2011). *nkx2.5*:Kaede-based lineage tracking has indicated that most of the ventricular myocardium is already condensed at the cardiac disc, but if all these cells then migrate with the emerging linear heart tube or stay behind has remained unresolved (Paffett-Lugassy et al., 2017). *nkx2.5*:Kaede further marks a seemingly distinct group of cells posterior and outside of the forming heart tube that also contributes myocardial progenitors to the distal ventricle and OFT (Zeng and Yelon, 2014); how these cells connect to the other ventricular progenitors remains to be uncovered. Position-based cell labeling experiments have mapped BA origins to the medio-central region of the heart-forming ALPM that corresponds to the expression domain of *nkx2.5* and *gata4* (Hami et al., 2011), with the proximal-most part of the BA arising from *nkx2.5* reporter-expressing pharyngeal arch 2 mesoderm (Paffett-Lugassy et al., 2017). Altogether, these analyses support a model of addition of the majority of late-differentiating myocardium to the ventricle and BA formation after establishment of the linear heart tube.

The T-box transcription factor Tbx1 is expressed within the CPF of various chordates and directs cardiac development by maintaining proliferation and suppressing differentiation of SHF cardiac progenitor cells (Chen et al., 2009; Diogo et al., 2015; Onimaru et al., 2011; Scambler, 2010). Impaired *TBX1* in humans results in DiGeorge Syndrome (Scambler, 2010) with variable cardiac defects including Tetralogy of Fallot, OFT defects, and an interrupted aortic arch, defects recapitulated by *Tbx1*-mutant mice (Jerome and Papaioannou, 2001; Lindsay et al., 2001; Merscher et al., 2001). The zebrafish *tbx1* mutant *van gogh* (*vgo*) displays among other phenotypes defects in the pharyngeal arches and a smaller BA, underlining the conserved function of Tbx1 in CPF control (Choudhry et al., 2013; Piotrowski et al., 2003). The 12.8 kilobases (kb) upstream of the murine *Tbx1* gene as transgenic reporter principally recapitulate endogenous expression through separable Forkhead factor-binding enhancers that drive pharyngeal/anterior endoderm versus mesoderm expression, including activity in the OFT (Hu et al., 2004; Maeda et al., 2006; Yamagishi et al., 2003). While these enhancers are sufficient in transgenic reporters, endogenous *Tbx1* expression is redundantly coordinated with additional activating and suppressing elements in the vicinity of the locus (Zhang and Baldini, 2010).

Here, we have isolated upstream *cis*-regulatory elements from the zebrafish *tbx1* locus to visualize the dynamics of ventricle and OFT formation. Combining selective plane illumination microscopy (SPIM) imaging with genetic and optogenetic lineage tracing, we captured the formation of the linear heart tube with concomitant migration of an undifferentiated sheath of *tbx1* reporter-expressing cells that are continuously added to, and gradually differentiate at, the arterial pole of the heart. Meanwhile, BA progenitors reside in the *tbx1* reporter-positive pharyngeal ALPM and migrate later towards the late-differentiating distal pole of the ventricle to become smooth muscle. Combining chemical and genetic perturbations, we found a distinct temporal requirement for FGF signaling in controlling ventricle and BA size versus BA specification. In contrast to models that postulate a distant cellular origin and stepwise addition of SHF cells, our findings establish that incorporation of the zebrafish ventricle, and OFT structures into the linear heart tube is a continuous process with distinct phases of FGF activity during CPF differentiation.

## Results

### *cis*-regulatory elements of the zebrafish *tbx1* locus are active in cardiac progenitors

In our ongoing efforts to isolate *cis*-regulatory elements active within the lateral plate mesoderm (LPM) to observe cardiovascular cell fate partitioning, we generated zebrafish *tbx1* reporter transgenics based on the high ranking of *tbx1* expression in transcriptome analysis of zebrafish LPM (within top-20 enriched genes) (Mosimann et al., 2015). Transgenic reporters in mice have established *5*’ core regulatory elements sufficient for recapitulating *Tbx1* expression (Yamagishi et al., 2003). Consistently, we observed specific EGFP reporter activity driven by the 3.2 kb upstream region of zebrafish *tbx1* in embryos carrying transgenic insertions of *Tg(-3.2tbx1:EGFP)^zh703^*, with minimal variability between six individual transgenic lines (Figure 1B, Supplementary Figure 1, Supplementary Table 1).

In late epiboly, tbx1:EGFP akin to endogenous *tbx1* expression broadly labels a dorsal/anterior domain (Figure 1C, Supplementary Figure 1). During somitogenesis, *tbx1*:EGFP expression is detectable in anterior bilateral domains (Figure 1D,E) and at 36 hpf in the pharyngeal arches and in the heart (Figure 1F). While we do not detect significant endogenous *tbx1* mRNA expression in the heart consistent with previous reports (Nevis et al., 2013), we readily observed *tbx1*:EGFP expression in cardiac precursors, indicating distinct dynamics of our reporter compared to endogenous *tbx1* akin to mouse *Tbx1* reporters (Supplementary Figure 1) (Hu et al., 2004; Maeda et al., 2006; Yamagishi et al., 2003). To resolve *tbx1* reporter expression in relation to the *drl*-labelled LPM (Mosimann et al., 2015), we performed *in toto* panoramic SPIM imaging (Schmid et al., 2013) on *tbx1*:EGFP;*drl*:mCherry transgenic embryos from gastrulation (70% epiboly) to mid-somitogenesis (15 hpf) (Supplementary Video 1). At late epiboly to tailbud stages, when the LPM condenses around the embryo margin, we detected overlapping *tbx1*:EGFP expression in *drl*:mCherry-expressing LPM cells, medial within the ALPM and lateral-most within the *tbx1* reporter-expressing domain (Figure 1G,H). These cells condensed further at the margin of the *tbx1*:EGFP-expressing domain throughout early somitogenesis (Figure 11,J). Thus, a sub-population of the *tbx1* reporter-expressing cells represent ALPM, while the broad medial field of *tbx1* reporter-positive cells potentially represent endoderm precursors, as we observe labeling of endodermal derivatives in i) 3 dpf *tbx1*:EGFP-expressing embryos, as well as ii) in CreERT2/lox-mediated lineage tracing of (*Tg(-3.2tbx1:creERT2)^zh703^* transgenics (Supplementary Figure 2). Moreover, cranial cartilage is labelled by *tbx1* reporter expression and with *tbx1:creERT2*-mediated lineage tracing, suggesting additional *tbx1* reporter expression in neural crest lineages (Supplementary Figure 2). Taken together, transgenic zebrafish reporter expression based on the upstream 3.2 kb *tbx1 cis*-regulatory region approximates key aspects of reported Tbx1 activity (Chen et al., 2009; Diogo et al., 2015; Garg et al., 2001) and visualizes a dynamic anterior endoderm and mesoderm domain that includes ALPM.

### ALPM-derived *tbx1* reporter-expressing cells contribute to both venous and arterial poles of the heart

To resolve cardiac *tbx1*:EGFP expression, we analyzed *tbx1:EGFP* transgenics co-stained for the differentiated cardiomyocyte-expressed Myosin Heavy Chain 1E (MYH1E, MHC) (Figure 2A). At 26 hpf, when the differentiating cardiomyocytes in the linear heart tube represent FHF derivatives (Mosimann et al., 2015; de Pater et al., 2009), we detected *tbx1* reporter expression in most of the differentiated ventricular cardiomyocytes and additionally in two MHC-negative domains at the IFT and OFT (Figure 2A). At 26 hpf, we detected *tbx1*:EGFP;Isl1 double-positive cells at the IFT of the linear heart tube (Figure 2B,C), consistent with inferred SHF identity of IFT cells (Caputo et al., 2015; de Pater et al., 2009; Witzel et al., 2017).

**Figure 2:**
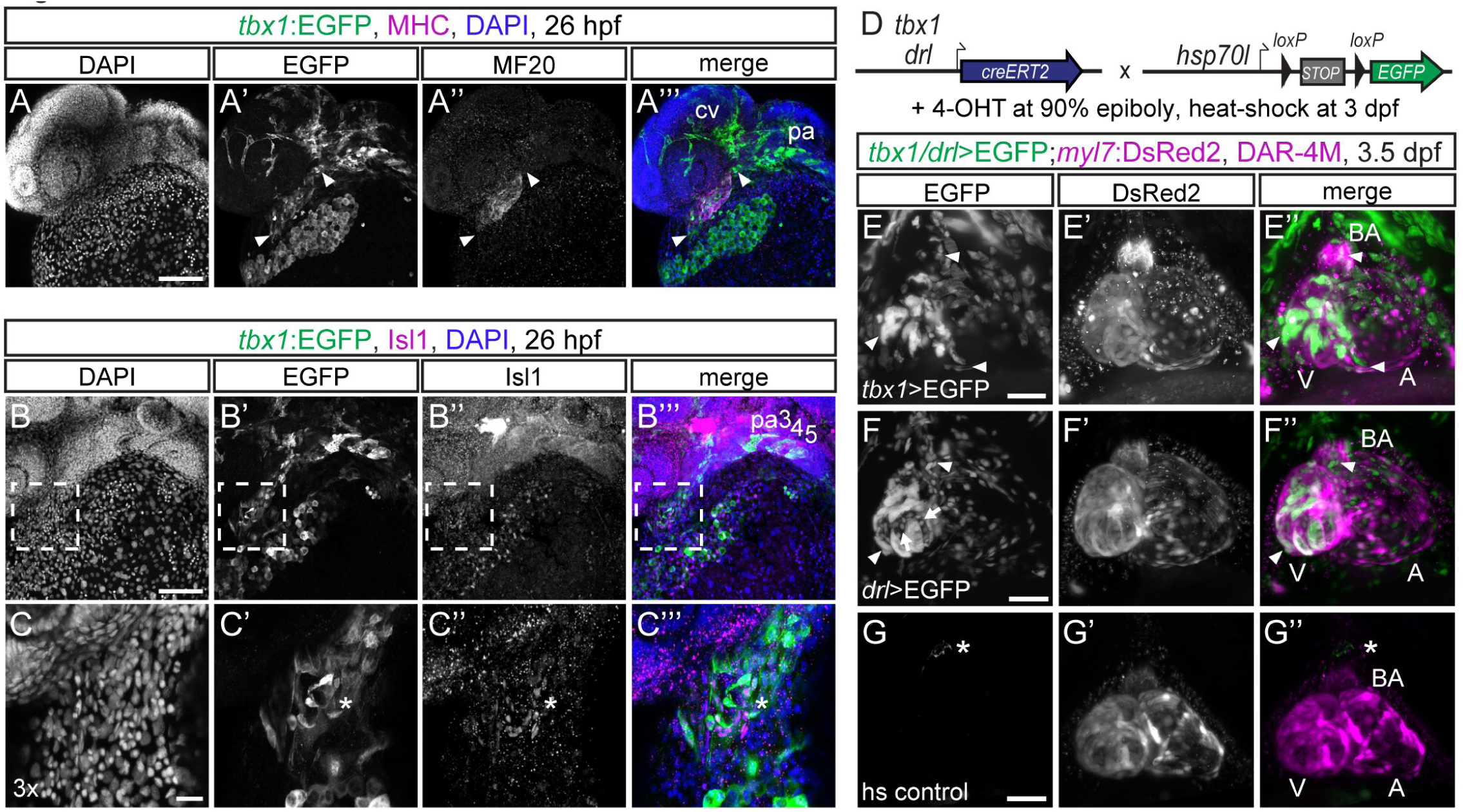
*tbx1* reporter-expressing cells contribute to LPM-derived cardiac lineages. (**A-C**) Maximum intensity projections of wholemount *tbx1*:EGFP-expressing embryos counterstained for anti-EGFP and anti-MHC (**A**) or anti-Isl1 (**B,C**) at 26 hpf; lateral views, anterior to the left. (**A**) *tbx1* reporter-expression can be detected in the MHC-positive linear heart tube and in the MHC-negative poles at the cardiac inflow and outflow tracts (arrowheads). *tbx1*:EGFP also marks the pharyngeal arches (pa) and endothelial cells of the cranial vasculature (cv). (**B,C**) *tbx1* reporter-expressing cells at the inflow tract co-express the SHF marker |s|1 (asterisks **C**); (**C**) depicts a 3x magnification of the framed area in (**B**). (**D**) Lineage tracing of *tbx1* and *drl* reporter-expressing cells. *tbx1:creERT2* or *drl:creERT2* transgenics, respectively, were crossed to the ubiquitous *hsp70l:Switch loxP* tracer line, 4-OHT-induced at 90% epiboly, and heat-shocked at 3 dpf. (**E-G**) Live SPIM imaging of still hearts of representative lineage-traced and control embryos; maximum intensity projections of ventral views, anterior to the top. (**E**) *tbx1*:creERT2 lineage tracing (*tbx1*>EGFP) at late gastrulation labels *myl7*:DsRed2-expressing cardiomyocytes in the ventricle (V) and inflow tract of the atrium (A), and DAR-4M-stained cells in the BA (arrowheads). (**F**) *drl*:creERT2-mediated lineage tracing (*drl*>EGFP) at 90% epiboly marks all cardiomyocytes in the ventricle and atrium, BA cells (arrowheads), and the endocardium (arrows). (**G**) *tbx1:creERT2;hsp70l:Switch* transgenics without 4-OHT treatment and heat-shocked at 3 dpf show no specific EGFP expression (asterisks marks auto-fluorescent pigment cell). Scale bars 100 μm.

To corroborate which cardiac lineages form from *tbx1* reporter-expressing cells, we performed genetic lineage tracing with *tbx1:creERT2* and *hsp70l:Switch (Tg(-1.5hsp70l:loxP-STOP-loxP-EGFP, cryaa:Venus)^zh701^* (Figure 2D). 4-OHT induction of CreERT2 at shield stage to 90% epiboly (6-9 hpf) labelled ventricular cardiomyocytes, including the distal ventricle and OFT region, scattered atrial cells around the IFT, and the Diaminorhodamine-4M AM (DAR-4M)-reactive smooth muscle cells in the BA (Grimes et al., 2006) (Figure 2E, Supplementary Figure 3). These cardiac descendants of *tbx1* reporter-expressing cells likely derive from ALPM: pan-LPM lineage tracing using *drl:creERT2* from 90% epiboly broadly marks atrial and ventricular myocardium plus the DAR-4M-stained BA (Figure 2F, Supplementary Figure 3), in line with previous LPM labeling by *drl:creERT2* (Felker et al., 2016; Henninger et al., 2016; Mosimann et al., 2015) and selective ALPM tracing of OFT structures (Guner-Ataman et al., 2013; Paffett-Lugassy et al., 2017). While *drl*-mediated ALPM lineage tracing labeled the endocardium, cranial vessels (Figure 2F, Supplementary Figure 3), and several head muscle groups (Supplementary Figure 3), *tbx1* reporter expression and lineage tracing labeled the same structures without endocardium, but additionally also craniofacial cartilage and endoderm derivatives, in line with Tbx1 as CPF marker (Diogo et al., 2015) (Figure 2A, Figures S2, S3). These results indicate that the *tbx1* reporter expression domain entails ALPM-derived cells that contribute to cardiac lineages including IFT and OFT structures (Figure 2E-G).

### *tbx1* reporter-expressing progenitors contribute to and migrate together with the linear heart tube

By end point analysis in zebrafish, different positions and migration dynamics have been assigned to IFT-, ventricle-, and OFT-contributing undifferentiated SHF progenitors, including delayed migration from pharyngeal ALPM (Hami et al., 2011; Lazic and Scott, 2011), condensation with the forming cardiac disc (Paffett-Lugassy et al., 2017), and localization posterior to the forming cardiac cone (Zeng and Yelon, 2014). To visualize the cardiac *tbx1* reporter-expressing cells, we SPIM-imaged *tbx1:EGFP;drl:mCherry* embryos from 16 hpf (14 ss) to 22-24 hpf to capture all stages of heart field migration up to the onset of heart beat (Supplementary Video 2). Of note, at these stages *drl*:mCherry expression in the heart field gradually confines from earlier pan-LPM expression to restricted expression in FHF-derived lineages, and robust expression in the endocardium (Mosimann et al., 2015). At 14 ss, the *drl*:mCherry-expressing heart field is arranged as a bilateral LPM domain as is the nearly overlapping *tbx1* reporter-expressing field before condensing at the midline (Figure 3A,B). From approximately 16 ss onwards, while midline-centered migration and formation of the *drl*:mCherry-positive cardiac disc completed, *tbx1*:EGFP-positive cells contributed to the cardiac disc (Figure 3B) and further disseminated from the bilateral *tbx1* reporter domain to migrate medially along *drl*-positive endothelial progenitors (Figure 3C). By 22 hpf, *tbx1* reporter-positive cells formed a sheath of cells at the prospective arterial pole of the both *tbx1* and *drl* reporter-expressing linear heart tube (Figure 3D); 3D segmentation confirmed that in its entirety, the *tbx1* reporter-positive cell population surrounded the growing endocardium like a sleeve trailing outwards to the still bilateral progenitors (Figure 3E,F, Supplementary Video 3). These observations suggest co-migration of *tbx1* reporter-expressing cells as part of the forming and jogging linear heart tube and continuous with the prospective arterial pole (Figure 3E,F).

**Figure 3:**
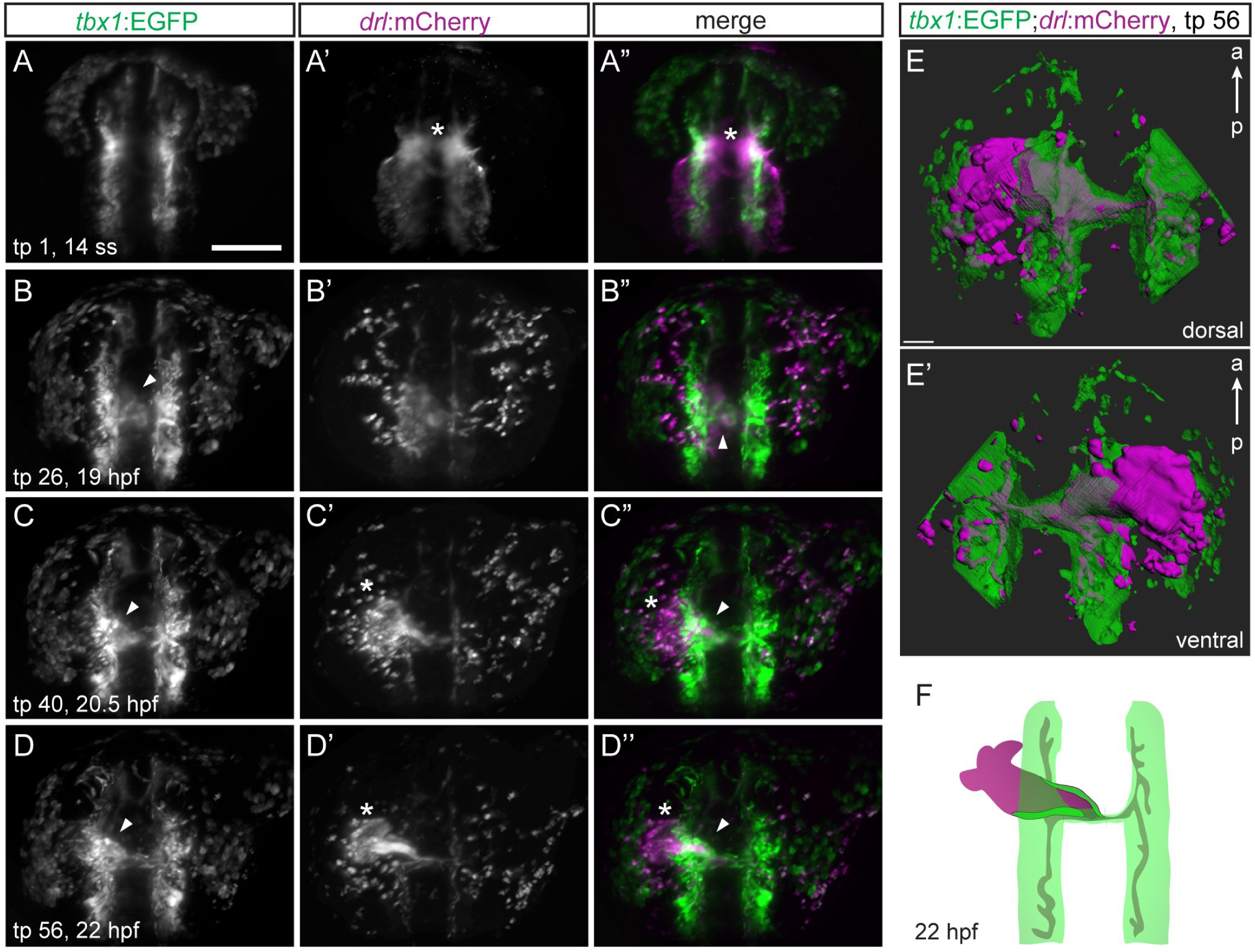
A *tbx1* reporter-expressing sheath forms at the base of the FHF-derived heart tube. (**A-C**) Maximum intensity projections of representative stages from SPIM-imaged *tbx1:EGFP;drl:mCherry* double-positive transgenic embryos; dorsal views, anterior to the top. Imaging was initiated at 14 ss (16 hpf) and cardiac development followed until LHT stage (22-23 hpf). (**A**) 14 ss stage embryo at onset of medial FHF migration (asterisk). (**B**) The forming cardiac disc already contains *tbx1*:EGFP-positive cells (arrowhead). (**C,D**) *tbx1*:EGFP-positive cells (arrowhead) assemble at the base of the extending *drl* reporter-expressing heart cone (asterisk in **C**) and are contained in the LHT (arrowhead in **D**), note absence of *tbx1*:EGFP reporter-expressing cells at the leading edge (asterisk in **D**) of the forming heart tube. (**E**) 3D segmentation (dorsal and ventral view) revealing a *tbx1* reporter-expressing sheath of cells engulfing the *drl* reporter-expressing endocardium at 22-23 hpf. (**F**) Schematic of *tbx1* and *drl* reporterexpressing cell arrangements at the end of imaging. Scale bars 50 μm (**E**), 200 μm (**A-D**).

### The late differentiating *tbx1* domain is continuous with the early forming myocardium

To understand the formation of the arterial pole of the heart tube in greater detail, we imaged *tbx1:EGFP;myl7:DsRed2* double-transgenic embryos from 18-30 hpf (Supplementary Video 4, Supplementary Figure 4). The slow folding of DsRed2 during this time frame discriminates between the early differentiated, early DsRed2-positive (FHF-assigned) and the later differentiating, DsRed2-negative (SHF-assigned) *myl7*-expressing cardiomyocytes (de Pater et al., 2009; Witzel et al., 2012). Correspondingly, we detected myl7:DsRed2 expression in the migrating and differentiating heart tube from 24 hpf onwards (Supplementary Figure 4). We observed *tbx1* reporter-expressing cells at the arterial pole of the heart tube i) that based on absence of *myl7*:DsRed2 expression throughout 24-30 hpf correspond to the later differentiating SHF myocardium, and ii) that were connected to, and migrated together with, the *myl7*-positive myocardium (Supplementary Figure 4).

To overcome the complications of live tracking associated with the onset of the heart beat at 24 hpf with continued ventral and rostral heart tube migration (Guner-Ataman et al., 2013; Hami et al., 2011; Lazic and Scott, 2011; Mosimann et al., 2015), we performed SPIM-based high-speed imaging and reconstruction of the beating zebrafish heart from 28 to 52 hpf (Mickoleit et al., 2014). We imaged *tbx1:EGFP;myl7:DsRed2* embryos from a lateral view (right side) to optimally resolve the migrating and looping ventricle (Figure 4A-C, Supplementary Video 5). Linking to our previous time course, we observed the *tbx1*:EGFP-positive/*myl7*:DsRed2-negative cells connected to the differentiating, *myl7*:DsRed2-fluorescent myocardium at 28 hpf (Figure 4A); displacement of these cells due to the beating differentiated myocardium supports their continuous incorporation into the heart tube (Supplementary Video 6). Throughout ventricle looping, the initially solely *tbx1* reporter-positive cells became re-arranged within the ventricle, intercalated with *myl7*:DsRed2-expressing differentiated cardiomyocytes, and gradually turned on *myl7*:DsRed2 expression (Figure 4B,C), in agreement with the previously described late SHF-derived myocardial differentiation (Lazic and Scott, 2011; de Pater et al., 2009; Zhou et al., 2011). Imaging of still hearts showed that all ventricle-incorporated tbx1:EGFP-expressing cells were differentiated and expressed *myl7*:DsRed2 by 54 hpf (Figure 4D). In addition, *tbx1* reporter-expressing undifferentiated cells that gradually upregulated *myl7* expression also appeared at the IFT (Figure 4A-C).

**Figure 4:**
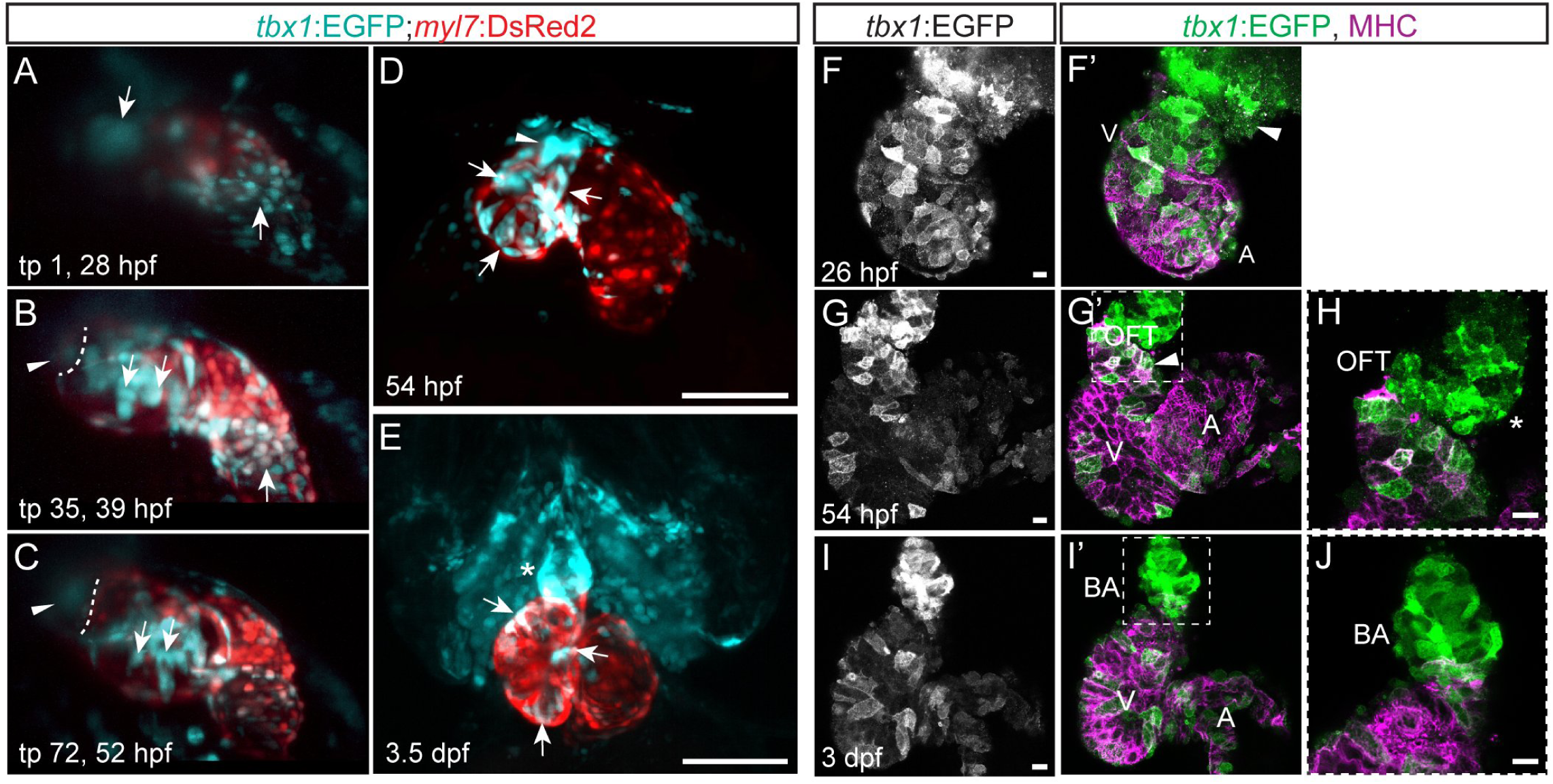
*tbx1* reporter-expressing SHF-derived myocardial precursors are connected to the FHF myocardium during heart tube stages. (**A-C**) Maximum intensity projections of representative stages of a high-resolution reconstruction of the beating heart of a *tbx1*:EGFP;*myl7*:DsRed2 double positive transgenic between 28-52 hpf; lateral view (right side) of the embryo, anterior to the top, ventricle to the upper left, atrium to the lower right, cardiac imaging phase 27. Arrows indicate *tbx1*+/*myl7*-cells at the OFT and IFT at the beginning of the time-lapse (**A**) that gradually turn on *myl7* expression (**B,C**). The dashed line (**B,C**) indicates the distal end of the ventricle and the arrowheads point to *tbx1*:EGFP-expressing cells at the OFT that never cross into the ventricle and are likely BA precursors. (**D,E**) Maximum intensity projections of SPIM-imaged *tbx1:EGFP;myl7:DsRed2* double-positive transgenic hearts stopped from contracting with BDM at 54 hpf or 3.5 dpf, respectively; ventral views, anterior to the top. *tbx1* reporter expression can be detected in differentiated *tbx1/myl7* reporter double-positive cardiomyocytes (arrows **D,E**), the *tbx1*+/*myl7*-OFT at 54 hpf (arrowhead **D**), and the *tbx1*+/*myl7*-BA at 3.5 dpf (asterisk **E**). (**F-J**) Top-down 2-μm confocal section of isolated zebrafish hearts at 26 (**F**), 54 (**G, H**) and 72 hpf (**I, J**) from tbx1:EGFP, counter-stained with anti-GFP and anti-MHC; OFT/BA to the right, atrium (A) to the top left, ventricle (V) to bottom left. (**F**) *tbx1*:EGFP is expressed at the MHC-negative arterial pole of the heart tube (arrowhead). (**G,H**) At 54 hpf, cardiomyocytes of the later-differentiated distal ventricle express *tbx1*:EGFP (arrowhead), as do MHC-negative progenitors of the OFT (asterisks). (**I,J**) The differentiated BA at 3 dpf is positive for *tbx1*:EGFP. Scale bars 10 μm (**F-J**), 200 μm (**D,E**).

In our high-speed time-lapse, we noted that from time point 35 (approx. 39 hpf) a new cluster of *tbx1*-reporter-expressing cells appeared at the OFT that expanded throughout the rest of recorded cardiac development, but never crossed into the ventricle (Figure 4B,C, Supplementary Video 5); we hypothesized that these cells are BA progenitors. Indeed, we detected tbx1:EGFP expression at 3-4 dpf in the differentiated BA smooth muscle (Figure 4E), consistent with our genetic lineage tracing (Figure 2E). We corroborated our live imaging data by examining *tbx1*:EGFP reporter expression in dissected hearts of transgenic embryos (Figure 4F-J); we observed *tbx1*:EGFP-positive cells at the distal ventricle of 26 hpf hearts that did not express MHC at this time (Figure 4F). At 54 hpf, we detected differentiated cardiomyocytes expressing *tbx1*:EGFP as well as potential BA precursors at the OFT that did not express MHC (Figure 4G,H). At 3 dpf, the differentiated smooth muscle but not the endothelium of the BA was clearly marked by tbx1:EGFP expression (Figure 4I,J, Supplementary Figure 4).

Altogether, our live imaging data confirmed by analysis in isolated hearts i) provide real-time observation of late-differentiating ventricle myocardium, and ii) confirm that *tbx1* reporter-positive cells that co-migrate with the early differentiating cardiac cone subsequently differentiate to ventricular cardiomyocytes, including the late-differentiating myocardial pool at the arterial pole (de Pater et al., 2009; Zhou et al., 2011). Moreover, we detect a second phase of *tbx1* reporter-expressing cells adding to the heart at later stages and potentially making the smooth muscle of the BA.

### The *tbx1* reporter-expressing sheath forms the majority of the ventricular myocardium

While corresponding to our *tbx1:creERT2* lineage tracing (Figure 2E), active EGFP reporter expression is not strict evidence for lineage association. To confirm the lineage contribution of the *tbx1* reporter-expressing cardiac sheath to the ventricular myocardium and OFT, we performed optogenetic lineage tracing using the transgenic line *Tg(-3.2tbx1:H2B-Dendra2)^zh704^* (subsequently as *tbx1:Dendra2*); in *tbx1*:Dendra2, a nuclear histone 2B-linked Dendra2 fluorophore is constitutively green-fluorescent and turns irreversibly red upon photoconversion (Gurskaya et al., 2006). We photoconverted tbx1:Dendra2-positive cells of the cardiac cone and trailing sheath at 22 hpf (Figure 5A) and detected Dendra2-red positive cardiomyocytes at 3.5 dpf throughout the ventricle, including the SHF-assigned distal portion and the inner curvature (n=10/11, N=4; Figure 5B, Supplementary Figure 5). These observations suggest that the *tbx1*:Dendra2-expressing sheath contributes to FHF- and SHF-linked ventricular myocytes. When converting tbx1:Dendra2-expressing cells at the base of and posterior to the cone in the area previously assigned to harbor additional myocardial SHF precursors (Zeng and Yelon, 2014) (Figure 5C), we could detect a few distal cardiomyocytes only on the dorsal side of the ventricle labelled by Dendra2-red (n=3/3, N=1; Figure 5D). We identified the majority of the ventricle to be derived from *tbx1*:Dendra2-expressing cells present already in the cone, and only a small fraction of cardiomyocytes accrued later. The SHF contribution was previously reported to comprise 40-50% of the ventricular myocardium (Lazic and Scott, 2011; Mosimann et al., 2015; Zhou et al., 2011); thus, our data suggests that the majority of late-differentiating cardiomyocyte progenitors enter the heart trailing the early differentiating ventricular progenitors by 22 hpf as part of a continuous *tbx1* reporter-expressing cell sheath.

**Figure 5:**
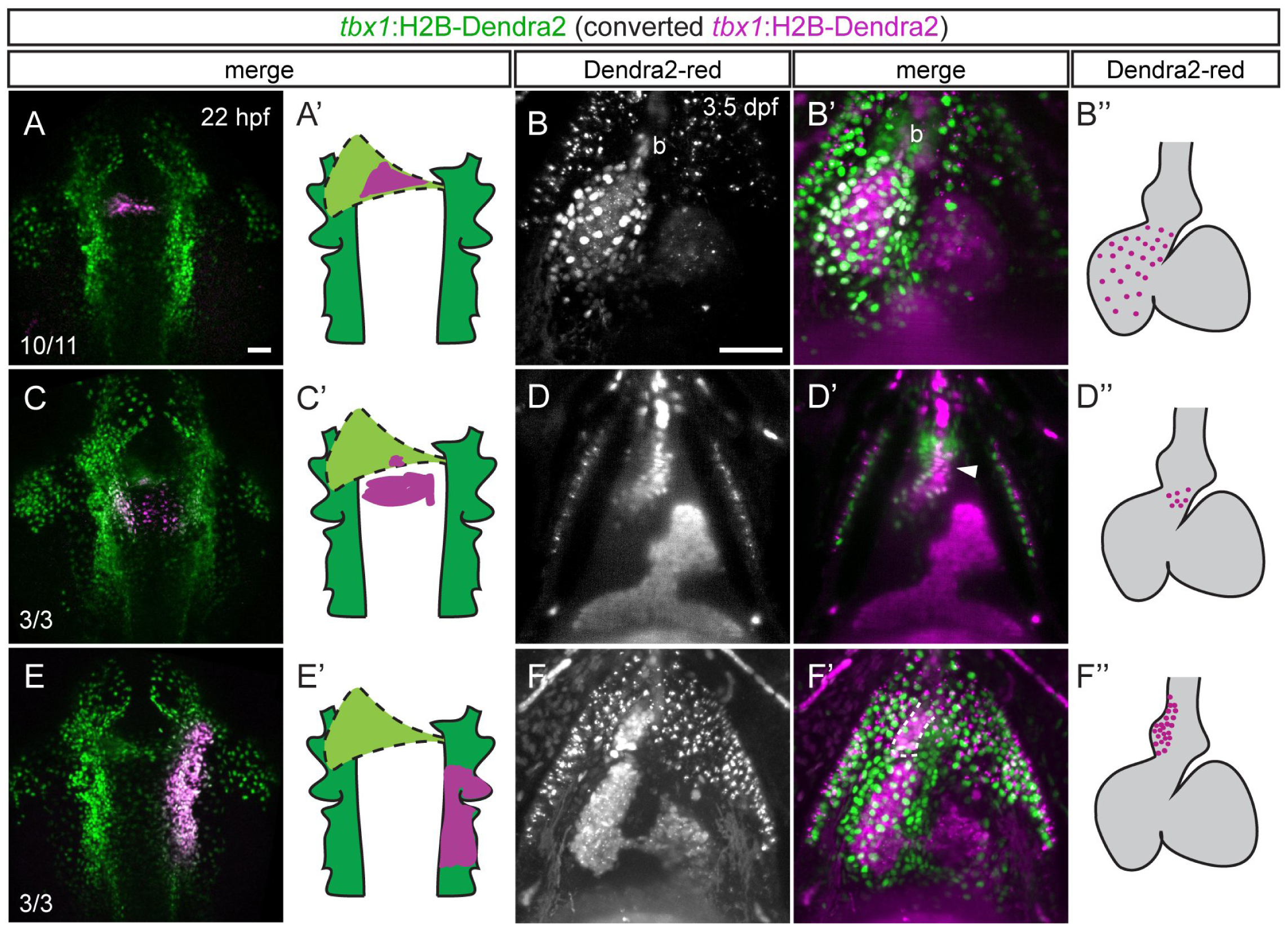
The *tbx1* reporter-expressing cone forms ventricular myocardium but not the smooth muscle-containing outflow tract. (**A,C,E**) Maximum intensity projections and schematics of representative photoconverted *tbx1*:Dendra2 embryos; dorsal views, anterior to the top. At 22 hpf, *tbx1*:Dendra2-expressing embryos were illuminated with UV light in a confined region of interest to convert Dendra2-green to Dendra2-red in specific *tbx1* reporter-expressing domains. (**B,D,F**) SPIM-imaged hearts and graphical representations of embryos photoconverted as in **A,C,E** and stopped from contracting with BDM at 3.5 dpf; maximum intensity projections (**B,F**) or optical Z-section (**D**), ventral views, anterior to the top. bcardiomyocytes marked by anti-MHC.(**A,B**) Dendra2-red-positive sheath cells give rise to ventricular cardiomyocytes, including the SHF-derived distal ventricle, but not to the BA. Red signal within the BA (asterisks) derive from autofluorescent blood also detected in non-photoconverted tbx1:Dendra2 embryos (Figure S6). (**C,D**) Medial migrating cells posterior to the cardiac cone contribute to the most distal myocardium at the dorsal side of the heart (arrowhead) and to a proximal portion of the BA in 3/3 analyzed embryos. (**E,F**) Photoconversion of a broad area in the *tbx1*:Dendra2-positive pharyngeal ALPM posterior to and on the right of the linear heart tube marks the right side of the BA (dotted outline). Red signals in the chambers, on the pericardium and within cranial vessels are due to unspecific autofluorescence also detected in controls (Supplementary Figure 6). Scale bars 50 μm.

In contrast to ventricular cardiomyocytes, we could not detect any Dendra2-red cells in the BA after photoconversion at the trailing end of the cardiac cone (n=0/11, N=4; Figure 5A,B, Supplementary Figure 5). In contrast, consistent with earlier position-based lineage tracing (Hami et al., 2011; Paffett-Lugassy et al., 2017), we detected Dendra2-red-positive cells in the BA when converting the pharyngeal ALPM lateral to the forming cardiac cone (n=15/15, N=4; Figure 5E,F). The right and left sides of the BA were exclusively formed from the corresponding side with no discernible cross-over (n=3/3 left side and n=3/3 right side, total n=6/6, N=3; Figure 5E,F, Supplementary Figure 6). Moreover, different regions of the pharyngeal LPM contributed to different parts of the BA on a proximal to distal axis and additionally labeled different craniofacial structures (n=3/3 proximal part, n=3/3 medial part, n=3/3 distal part, N=2; Figure 5E,F; Supplementary Figure 6). Taken together, our data document that the majority of, if not all, ventricular cardiomyocytes stem from a *tbx1* reporter-expressing progenitor sheath that participates in cardiac ALPM fusion, contributes to the linear heart tube, and trails the prospective arterial pole. In contrast, SHF-assigned BA precursors within the pharyngeal ALPM add to the heart at a subsequent stage as more distantly trailing cells, as has been reported for OFT lineages in mammals (Kelly et al., 2001; Mjaatvedt et al., 2001).

### FGF signaling selectively controls *tbx1+* progenitor addition to the zebrafish heart

FGF signaling influences cardiac patterning including SHF development in various chordates (Rochais et al., 2009). Correspondingly, zebrafish embryos mutant for *fgf8a* (*acerebellar, ace*) or upon *fgf8a* morpholino knockdown form a severely hypoplastic ventricle with comparably normal atrium (de Pater et al., 2009; Reifers et al., 2000). Moreover, chemical perturbations using the pan-FGF signaling inhibitor SU5402 (Marques et al., 2008; Mohammadi et al., 1997; Simoes et al., 2011) from earliest phases of heart formation result in reduced late-differentiating myocardium and loss of SHF marker expression at the arterial pole (Lazic and Scott, 2011; de Pater et al., 2009).

To determine the temporal requirement for FGF signaling on the different phases of cardiac contribution from the *tbx1* reporter-expressing ALPM field, we first revisited the *fgf8a* phenotype using the verified *fgf8a* translation-blocking morpholino MO3-*fgf8a*^ATG^ (Scholpp and Brand, 2001). We measured ventricle and BA size upon *fgf8a* perturbation in isolated hearts: *fgf8a* morphants still faithfully expressed tbx1:EGFP and formed a BA (Figure 6A-D), yet both ventricle and BA were significantly smaller (Figure 6E,F) consistent with previous reports (Marques et al., 2008; de Pater et al., 2009; Reifers et al., 2000). The impact on ventricle and BA size is unlikely from a strong disruption of FHF or SHF territories, as at 54 hpf expression of *drl*:EGFP that demarcates FHF-derived myocardium appeared insignificantly affected upon *fgf8a* perturbation (Figure 6G-I).

**Figure 6:**
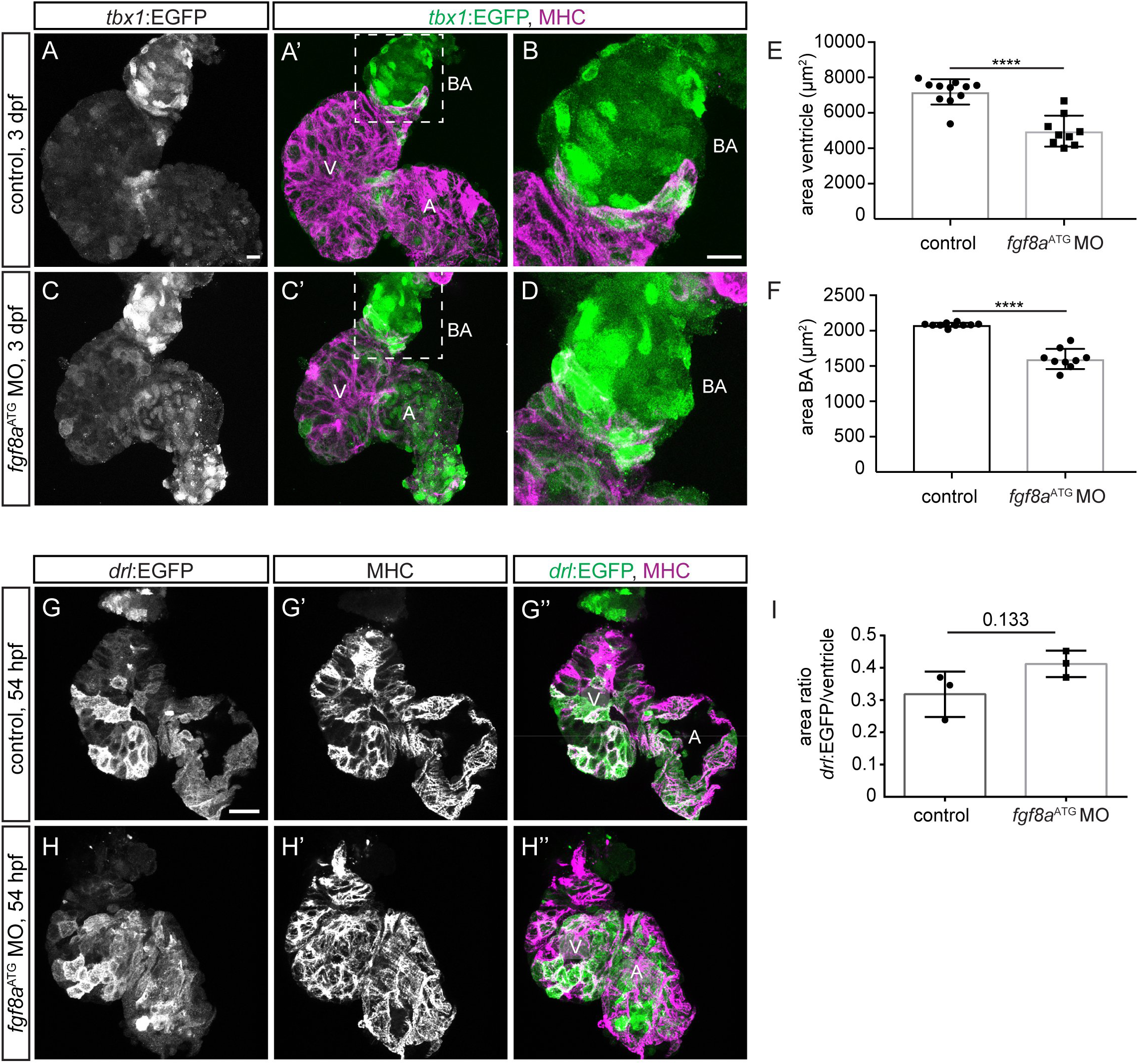
*fgf8a* knock-down leads to perturbed ventricle and BA formation. (**A-D**) Top-down 7 μm confocal section of uninjected control (**A,B**) and *fgf8a*^ATG^ morphant (**C,D**) hearts at 3 dpf. *tbx1*:EGFP, counter-stained with anti-GFP labels the bulbus arteriosus (BA) (**B** inset of **A’**; **D** inset of **C’**) and scattered ventricular (V) and atrial (A) cardiomyocytes, counter-stained with anti-MHC. (**E**) Quantification of the area of the ventricle in control hearts (n=11) compared to *fgf8a*^ATG^ morphant hearts (n=9) reveals significantly smaller ventricle upon knockdown of *fgf8a*. Each data point stands for the averaged ventricular area from one heart. (**F**) Quantification of the area of the BA in uninjected control hearts (n=10) compared to fgf8a^ATG^ morphant hearts (n=9) displays significantly smaller BA in the loss of *fgf8a*. Each data point represents the measurement of the BA area from one heart. (**G,H**) Top-down 3 μm confocal section of uninjected control (**G**) and *fgf8a*^ATG^ morphant (**H**) hearts at 54 hpf.*drl*: EGFP, counter-stained with anti-GFP labels FHF-derived cardiomyocytes marked by anti-MHC. (**I**) The ratio between the area of *drl*:EGFP positive cells and the area of the entire ventricle is not significantly different comparing controls and *fgf8a*^ATG^ morphants (P=0.133). Each data point presents calculated ratio from one heart. Means±s.d.. ****P<0.0001, unpaired t-test with Welch correction. Scale bars 10 μm.

To resolve the temporal influence of FGF signaling, we treated *tbx1:EGFP* embryos with SU5402 when *tbx1* reporter-expressing ventricular progenitors migrate to contribute to the forming heart tube. When we initiated SU5402 treatment before medial migration at 14 ss and perturbed FGF signaling throughout sheath migration (pulse treatment from 14 ss to 22 hpf), 5 μM SU5402-treated embryos showed diminished but still ongoing migration of the *tbx1*:EGFP-expressing sheath by 26 hpf (Figure 7A,B). In line with previous findings (Marques et al., 2008), at 3 days post-fertilization (dpf) the ventricle of these embryos appeared smaller, but nonetheless contained *tbx1*:EGFP-expressing cardiomyocytes (Figure 7C,D). We observed similar results when continuously perturbing FGF signaling with lower dose of 2 μM SU5402 from 14 ss onwards till imaging at 3 dpf (Figure 7E). These results demonstrate an impact of FGF signaling during *tbx1* reporter-expressing sheath migration, consistent with an early role of FGF signaling in ventricle formation.

**Figure 7:**
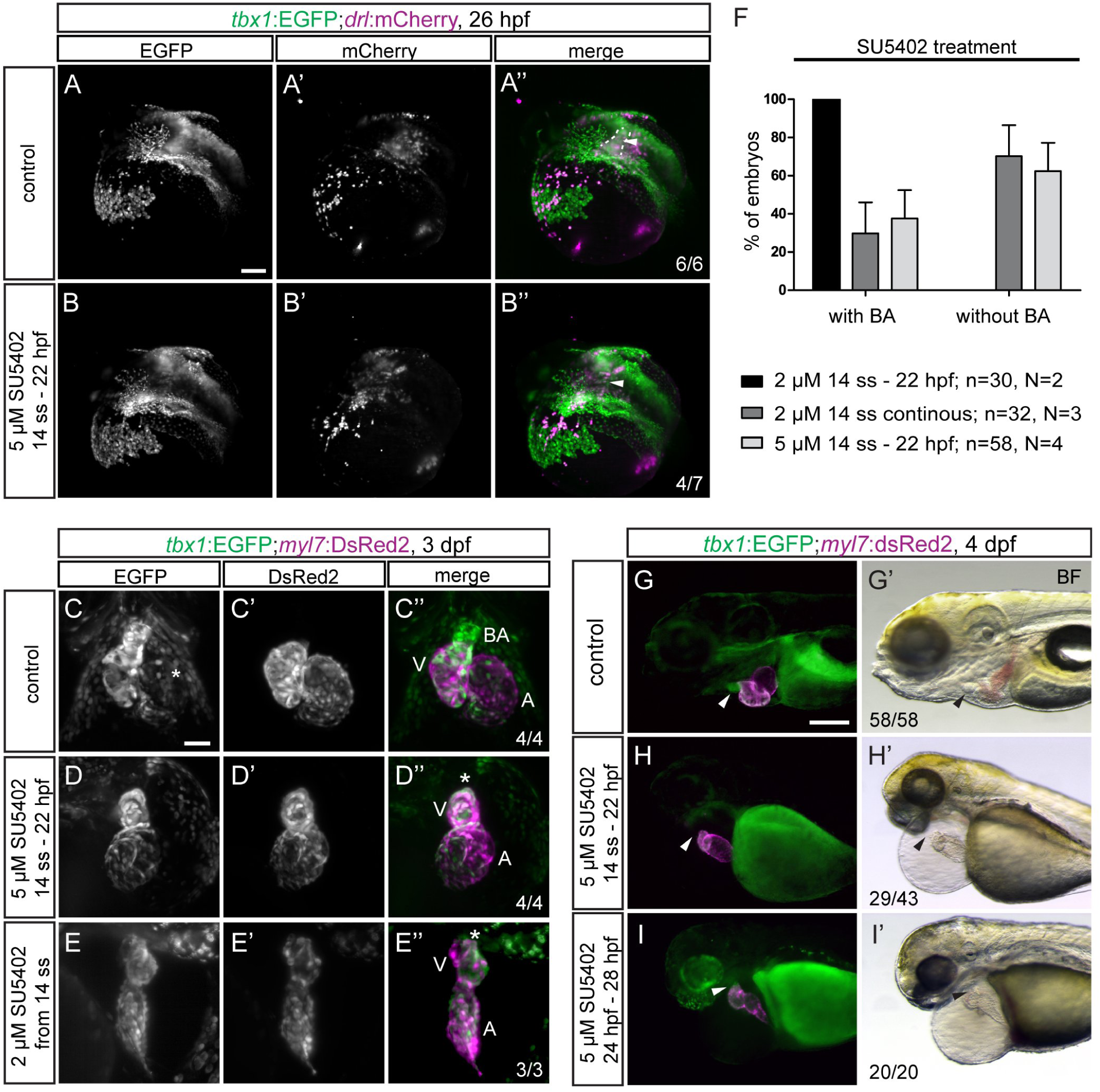
FGF signaling differentially affects *tbx1* reporter-expressing ventricular and bulbus arteriosus precursors during heart tube formation. (**A,B**) Maximum intensity projections of representative tbx1:EGFP;drl:mCherry transgenic controls or embryos treated with 5 μM SU5402 from 14 ss to 22 hpf; lateral/dorsal view, anterior to the left. FGF-signaling perturbed embryos show a defect in the tbx1:EGFP-expressing sheath (arrowhead) at the base of the forming heart tube (outline). (**C-E**) FGF-perturbed embryos retain normal contribution of *tbx1*:EGFP-expressing cardiomyocytes to the ventricle upon pulsed or continuous signaling inhibition; ventral views, anterior to the top. Instead, FGF signaling perturbation leads to a diminished ventricle size and abolishes addition of the *tbx1* reporter-expressing BA to the heart (asterisks). (**F-I**) SU5402 treatments affect BA development in a time- and concentration-dependent manner; n indicates the number of individual embryos analyzed per condition, N indicates the number of individual experiments performed. (**F**) Quantification of the concentration-dependent effect on BA formation in FGF signaling perturbed *tbx1*:EGFP;*myl7*:Red or DAR-4M-stained myl7:EGFP transgenics (see additionally Figure S7). (**G-I**) *tbx1*:EGFP;*myl7*:DsRed transgenic controls and embryos treated with DMSO or 5 μM SU5402 during (14 ss -22hpf) or after (24 hpf - 28 hpf) heart tube formation; lateral views, anterior to the left. Absent BA formation can only be observed in embryos treated with SU5402 from mid-somitogenesis to heart tube stages (arrow head **H**), but not when signaling inhibition is initiated at 24 hpf (arrow head **I**). Scale bars 50 μm (**C-E**), 100 μm (**A,B, G-J**).

In contrast to the mere reduction in ventricle size upon FGF inhibition, we observed a more striking and concentration-dependent effect on BA addition (Figure 7C-H). Even though BA addition takes place after primary heart tube formation at 24 hpf, BA formation was completely absent in a substantial number of embryos pulse-treated between 14 ss to 22 hpf with 5 μM SU5402 when assessed at 3 dpf as apparent by *tbx1*:EGFP or DAR-4M labelling (Figure 7D,F,H; Supplementary Figure 7; no BA in n=29/43 by *tbx1* reporter and n=5/15 by DAR-4M, total n=34/58, N=4; DMSO-treated controls: normal BA in n=58/58 by *tbx1* reporter and n=10/10 by DAR-4M, total n=68/68, N=4). While pulsing SU5402 treatment between 14 ss to 22 hpf with 2 μM SU5402 led to detectable BA differentiation (Figure 7F; present BA in n=15/15 by *tbx1* reporter and n=15/15 by DAR-4M, total n=30/30, N=2), continuously treated embryos from 14 ss onwards with 2 μM SU5402 lacked BA (Figure 7E,F; no BA development in n=4/6 by *tbx1* reporter and n=21/26 by DAR-4M, total n=25/32, N=3). In line, 5 μM SU5402 pulse-treatment from 24 hpf to 28 hpf or 34 hpf merely caused formation of smaller BA and never caused loss of the differentiated BA (pulse 24-28 hpf: n=20/20 by *tbx1* reporter, pulse 24-34 hpf: n=20/20 by DAR-4M; total n=40/40, N=3) (Figure 7I, Supplementary Figure 7).

Taken together, our data determine a temporally defined requirement for FGF signaling during formation and migration of the ventricle progenitor sheath. Further, our findings support a model in which differential levels and timing of FGF control the continuous addition of ventricle and BA progenitors to the heart tube, with a sensitive window for BA determination before its progenitors leave the pharyngeal ALPM.

## Discussion

The concept of distinct phases of differentiating cell types contributing to individual parts of the multi-chambered heart is deeply rooted in chordate evolution. Here, we visualized the formation of the zebrafish ventricle and OFT structures from a continuously differentiating progenitor sheath that emerges from *tbx1* reporter-expressing ALPM and surrounds the emerging endocardium. Our data connects, is consistent with, and extends previous end-point analyses of early-versus late-differentiating myocardium and the formation of OFT structures including the BA. We further reveal a temporal influence and sensitivity of FGF signaling on ventricle myocardium versus BA smooth muscle formation in zebrafish. Our work emphasizes cardiac development as part of a *tbx1*-expressing cardiopharyngeal progenitor field within the bilateral ALPM that is already significantly pre-patterned before its medial migration forms the heart.

Reporter expression and lineage tracing (Figures 1, 2, Figures S2, S3) established the isolated zebrafish *tbx1 cis*-regulatory region as a putative marker of the CPF that resides within the ALPM aside cranial endoderm and neural crest lineages. We have combined different strategies to demonstrate that our *tbx1* regulatory elements visualize early as well as late-differentiating cardiac lineages. When live-imaging *tbx1*:EGFP-expressing embryos from stages before heart field migration (14 ss) up to cardiac looping stages (54 hpf), we robustly detected *tbx1* reporter-expressing cells at the arterial pole of the ventricle that were connected to the differentiated *myl7*-expressing myocardium and started moving in sync with heartbeat of the primitive heart tube (Figures 3,4, Supplementary Videos 2-6); this observation suggests that undifferentiated *tbx1* reporter-expressing cells are already physically linked to beating cardiomyocytes. These *tbx1*:EGFP-expressing progenitors then gradually upregulated *myl7* expression, confirming their identity as cardiomyocyte progenitors equivalent to late-differentiating SHF-liked cells (Figure 4A-C, Supplementary Video 5) (de Pater et al., 2009; Witzel et al., 2012). Independent of continued *tbx1* reporter activity, genetic and optogenetic lineage tracking supports the notion that most, and likely all, FHF- and SHF-assigned ventricular cardiomyocytes are contained within the *tbx1* reporter-expressing sheath that contributes to the cardiac cone and trails into the bilateral ALPM (Figure 5A-D, Supplementary Figure 5). Of note, the SHF in mouse has been characterized as epithelial sheet that undergoes epithelial-to-mesenchymal transitions and tension changes during its addition to the heart tube (Francou et al., 2017). Our observations hint at similar dynamics of SHF cells in teleosts.

The observed addition of ventricle progenitors as continuous process connects previously observed key time points of SHF-assigned, late-differentiating myocardium addition to the zebrafish heart tube (Lazic and Scott, 2011; Mosimann et al., 2015; Paffett-Lugassy et al., 2017; Zhou et al., 2011). Nonetheless, the position of SHF-assigned cells during heart tube assembly had previously remained ambiguous. *nkx2.5:Kaede* reporter-based optogenetic lineage tracking showed that most if not all ventricular myocardium is already condensed at the cardiac disc, but if these cells then migrate with the emerging primitive heart tube or stay behind had remained unresolved (Paffett-Lugassy et al., 2017). A distinct group of *nkx2.5*:Kaede-expressing cells were found more posterior, seemingly outside of the forming heart tube, and shown to form a small portion of the distal ventricle and OFT myocardium (Zeng and Yelon, 2014). Our *tbx1* reporter transgenics now consolidate these data points as parts of the continuous cell sheath forming the ventricle.

On the contrary, smooth muscle cells in the BA were contributed from the still bilateral pharyngeal ALPM lateral and posterior to the developing heart tube (Figure 5E,F, Supplementary Figure 6), a development that also appears in our live imaging data (from approximately 39 hpf, time point 35 in Figure 4B,C, Supplementary Video 5). Inferred by reporter expression and not strictly lineage data, our analysis of dissected hearts at 54 hpf (Figure 4F-J) confirms the existence of a *tbx1* reporter-expressing, non-myocardial cell population at the OFT pole, the position where we later find the differentiated BA (Figure 4E,J). Addition of BA precursors to the distal ventricle has been reported as early as 48 hpf by 4,5-diaminofluorescein diacetate (DAF-2DA) staining that senses nitric oxide accumulating in the BA and likely requires functional maturation of smooth muscle (Grimes et al., 2006). A concise working definition (proposed in Figure 1A) to distinguish between distal ventricle myocardium, collagenous OFT myocardium of the CA, and BA could further consolidate the various reports of distinct waves of SHF-assigned cells to the heart tube that have been referred to by mixed nomenclature (Grimes and Kirby, 2009). Our data suggest a continuous addition of ventricle myocardium and a subsequent addition of smooth muscle progenitors as distinct phases of a continuous process.

FGF signaling has been previously reported to govern progenitor addition to the arterial pole of the zebrafish heart (Lazic and Scott, 2011; de Pater et al., 2009). Consistent with these previous findings, we observed a diminished ventricle upon pan-FGF signaling perturbation during *tbx1* reporter-expressing sheath formation (Figure 7C-E). We further detected absent BA formation when initiating chemical FGF signaling block already before cardiac cone formation (Figure 7F-I). In contrast, knockdown of the key cardiac FGF ligand gene *fgf8a* merely caused a smaller ventricle and BA (Figure 6A-F), akin to pulsed exposure to a low dose of SU5402 during ventricle formation and BA accrual at 14 ss-22 hpf (Figure 7F). To our knowledge, the drastic temporal influence of FGF signaling on BA formation has not been previously described in zebrafish. *Fgf8* depletion in mouse results in aberrant OFT development including severe aortic arch defects, a structure probably most comparable to the teleost BA (Abu-Issa et al., 2002; Frank et al., 2002). We only observed complete failure of BA formation when all FGF signaling was blocked before cardiac cone stages, but not when perturbed later during heart tube stages or by sub-penetrant doses of SU5402 (Figure 7F-I).

Our data is consistent with two effects of FGF signaling on zebrafish heart development after initial cardiac specification: first, FGF signaling regulates ventricular myocardium formation during medial migration of ventricular progenitors and cardiac cone formation; second, FGF signaling controls smooth muscle precursors residing in the pharyngeal ALPM prior to their OFT addition. Work on chick SHF explants has established that loss of FGF signaling blocks proliferation and causes myocardial differentiation, while elevating FGF signaling drives SHF cells into smooth muscle fates (Hutson et al., 2010). In *Ciona,* an FGF-driven regulatory circuit controls key cardiopharyngeal transcription factors including Tbx1/10, and regulates cell cycle dynamics to permit differentiation of individual cardiac lineages (Davidson et al., 2006; Razy-Krajka et al., 2017; Wang et al., 2017). These results are in line with our temporal and dose requirement for FGF during zebrafish ventricle and OFT formation (Figure 7), suggesting that BA progenitors already have an assigned fate during heart cone formation. Regulation of OFT progenitors while they reside in the pharyngeal LPM points towards an FGF activity gradient mediated by adjacent structures.

Of note, we also detected a late differentiating cardiomyocyte population at the venous pole, the IFT, of the heart (Figure 4A-C), in accordance with previous findings of late IFT myocardial differentiation (de Pater et al., 2009). *tbx1* reporter-expressing cells at the IFT concomitantly expressed Isl1, confirming their SHF signature (Figure 2B,C). We did not detect any Isl1 expression at the OFT, in accordance with other studies in zebrafish implicating Isl1 only in IFT development (Caputo et al., 2015; de Pater et al., 2009; Witzel et al., 2017). While we here focused on the development of the arterial pole, these observations warrant application of our *tbx1* reporter transgenics for the elucidation of zebrafish IFT formation.

Altogether, our data provide new insights into the dynamics of ventricle and OFT formation and integrate their mechanistic separation as distinct phases of a continuous developmental process in zebrafish.

## Methods

### Animal husbandry

Zebrafish (*Danio rerio*) were maintained, collected, and staged principally as described (Kimmel et al., 1995; Westerfield, 2007) and in agreement with procedures mandated by the veterinary office of UZH and the Canton of Zürich. Embryos were raised in temperature-controlled incubators without light cycle at 28°C unless specified differently in the text.

### Vectors and transgenic lines

All transgenic lines newly generated in this work have been assigned unique ZFIN designations. The upstream cis-regulatory region of the zebrafish *tbx1* gene (*ZDB-GENE-030805-5*) was amplified from zebrafish wildtype genomic DNA with primers *5’-GCTTATACGCACGACTGC-3’* (forward) and *5’-TGTGTCGATCGCGTATCGC-3’* (reverse) with the Expand Hi-Fidelity PCR kit (Roche). The 3242 bp upstream region of *tbx1* was TOPO-cloned into the *pENTR*^™^ *5’-TOPO*^®^ *TA Cloning*^®^ plasmid (Cat#59120; Invitrogen) according to manufacturer’s instructions to obtain *pAF006* (*pENTR/5_tbx1*).

Subsequent cloning reactions were performed with the Multisite Gateway system with LR Clonase II Plus (Cat#12538120; Life Technologies) according to the manufacturer’s instructions.

*tbx1:EGFP* (*pAF008* or *pDestTol2pA2_tbx1:EGFP*) and *tbx1:H2B-Dendra2* (*pAF048* or *pDestTol2pA2_tbx1:H2B-Dendra2*) were assembled from *pAF006* together with Tol2kit #*383* (*pME-EGFP*) or *pKP003* (*pME-H2B-Dendra2*), *#302* (*p3E_SV40polyA*), and *#394* (*pDestTol2A2*) as backbone (Kwan et al., 2007). This vector was used to generate transgenic strain *Tg(-3.2tbx1:EGFP)^zh702^* based in founder line I, and the additional lines for comparison as depicted in Figure S1 to ensure faithful transgene expression.

We cloned *tbx1:creERT2* (*pAF038* or *pDestTol2CY_tbx1:creERT2, alpha-crystallin:Venus*) by combining *pAF006* with *pCM293* (*pENTR/D_creERT2*) (Mosimann et al., 2011), Tol2kit vector *#302* (Kwan et al., 2007), and *pCM326*(*pDestTol2CY*, containing the *alpha-crystallin:Venus* cassette as transgenesis marker) as backbone(Mosimann et al., 2015). This vector was used to generate transgenic strain *Tg(-3.2tbx1:creERT2; cryaa:Venus)^zh703^*, which we selected as best line after screening several transmitting founders.

The *hsp70l:Switch* (*pAF040* or *pDestTol2CY_hsp70l:loxP-STOP-loxP-EGFP, alpha-crystallin:Venus*) transgene was assembled from *pDH083* (Hesselson et al., 2009) by transfer of the *loxP* cassette into *pENTR5’* (generating *pENTR/5’_hsp70l:loxP-STOP-loxP*), Tol2kit *#383* and *#302*(Kwan et al., 2007), and *pCM326* as backbone. This vector was used to generate transgenic strain *Tg(-1.5hsp70l:loxP-STOP-loxP-EGFP, cryaa: Venus)^zh703^*.

For Tol2-mediated zebrafish transgenesis, 25 ng/μL *Tol2* mRNA were injected with 25 ng/μL plasmid DNA (Felker and Mosimann, 2016; Kwan et al., 2007). F0 founders were screened for specific EGFP or *alpha-crystallin:YFP* expression, raised to adulthood, and screened for germline transmission. Single-insertion transgenic strains were established and verified through screening for a 50% germline transmission rate in outcrosses in the subsequent generations as per our previously outlined procedures (Felker and Mosimann, 2016).

Additional already established transgenic lines used in this study included *drl:mCherry* (*Tg(-6.3drl:mCherry)^zh705^)*), *myl7:DsRed2* (Chiavacci et al., 2012), *ubi:Switch* (Mosimann et al., 2011), and *drl:creERT2* (Mosimann et al., 2015).

### Morpholino injections

The *fgf8a*^ATG^ morpholino (MO3-*fgf8a*^ATG^: 5′-*GAGTCTCATGTTTATAGCCTCAGTA*-3′; ZFIN ID: *ZDB-MRPHLN0-050714-1*) that blocks translation (Scholpp and Brand, 2001) was obtained by Gene Tools, LLC and injected in the yolk of one-to four cell-stage embryos.

### Agarose sections

Transverse sections of agarose embedded embryos at 3 dpf were made as previously described (Gays et al., 2017). Briefly, embryos were fixed in 4% PFA, embedded in 6% low-melting point agarose (Sigma), cut into 130 mm vibratome sections (VT1000S, Leica), and mounted with DAPI-containing Vectashield (Cat#H-1200; Vector Laboratories).

### Whole-mount in situ hybridization

First-strand complementary DNA (cDNA) was generated from wildtype zebrafish RNA isolated with Superscript III First-Strand Synthesis kit (Cat#18080051; Invitrogen). DNA templates were generated using first-strand cDNA as PCR template and following primers: *EGFP* with 5′-*ATGGTGAGCAAGGGCGAGGAGC*-3′ (forward) and 5′-*TAATACGACTCACTATAGGG*-3′ (reverse); tbxl with 5′-*TATTCCGGATCCAACTCAGC*-3′ (forward) and 5′-*TTATCTGGGTCCGTAGTC*-3′ (reverse). For *in vitro* transcription, the T7 RNA polymerase promoter 5′-*TAATACGACTCACTATAGGG*-3′ was added to the 5′-end of reverse primers. *In situ* hybridization probes were made by *in vitro* transcription using T7 RNA polymerase and DIG-labeled NTPs (Cat#11277073910; Roche). RNA was precipitated with lithium chloride in ethanol and dissolved in DEPC water. Embryos were fixed in 4% Paraformaldehyde (PFA) overnight at 4°C, transferred into 100% methanol, and stored at -20°C until *in situ* hybridization. *In situ* hybridization of whole-mount zebrafish embryos was performed according to published protocols (Thisse and Thisse, 2008).

### Antibody staining

Embryos were fixed in 4% Formaldehyde, 0.1% TritonX in PEM (0.1 M PIPES, 2 mM MgSO4, 1 mM EDTA) for 2-4h at room temperature, washed in 0.1% PBS TritonX (PBSTx) and permeabilized in 0.5% PBSTx. Hearts from 26, 54 and 72 hpf zebrafish embryos were dissected in Tyrode’s solution (136 mM NaCl, 5.4 mM KCl, 1 mM MgCl_2_ x 6H_2_O, 5 mM D(+)Glucose, 10 mM HEPES, 0.3 mM Na_2_HPO_4_ x 2 H_2_O, 1.8 mM CaCl_2_ x 2H_2_O; pH 7.4) with 20 mg/mL BSA and fixed with Shandon^™^ Glyo-Fixx^™^ (Cat#9990920; Thermo Fisher Scientific^™^) for 20 min at room temperature. Blocking was done in blocking buffer containing 5% goat serum, 5% BSA, 20 mM MgCl_2_ in PBS and embryos/hearts incubated with primary antibodies diluted in blocking buffer at 4°C overnight. Primary antibodies used were anti-MHC (MF20 Supernatant, DSHB, 1:50), anti-GFP (Abcam, ab13970, 1:500 or Sigma, G1544, 1:100), and anti-Isl1 (GeneTex, GTX128201, 1:50). Alexa-conjugated secondary antibodies (A11039, A11004, A11008, and A11012 ThermoFisher Scientific) were added at 1:500 in 0.1% PBSTx at 4°C overnight. Embryos were washed several times in 0.1% PBSTx. DAPI-containing Vectashield (Cat#H-1200; Vector Laboratories) was added and embryos kept in the mounting medium until imaging. Before imaging, embryos were mounted in 1% low-melting point agarose. Dissected hearts were washed overnight in blocking buffer and mounted in the ProLong Gold antifade reagent with DAPI (Cat#P36935; ThermoFisher Scientific^™^).

### CreERT2-based lineage tracing

Lineage tracing experiments were performed by crossing female *hsp70l:Switch* or *ubi:Switch* reporter carriers with male *creERT2* driver transgenics (Felker and Mosimann, 2016). Embryos were induced using 4-OHT (Cat#H7904: Sigma) from fresh and/or pre-heated (65°C for 10 minutes) stock solutions in DMSO with a final concentration of 10 μM in E3 embryo medium as per our established protocols (Felker et al., 2016). Heat-shocks were performed for 60 min at 37°C in glass tubes in a water bath.

### Microscopy and image analysis

Stereo microscopy images were obtained on a Leica M205FA equipped with a Leica DFC450C digital camera. Confocal images of transverse sections were obtained on an inverted Zeiss LSM710 confocal microscope with a Plan-Apochromat 40x/1.3 Oil DIC M27 objective. Confocal imaging of whole-mounts and dissected hearts was done with a Leica SP8 upright confocal microscope using a HC PL APO 20x/0.5 Water objective and Leica SP8 inverted confocal microscope using a HC PL APO CS2 63x glycerol/NA 1.3 objective, respectively.

SPIM/lightsheet microscopy was performed on a Zeiss Z.1. Embryos were embedded in 1% low-melting point agarose and 0.016% Ethyl 3-aminobenzoate methanesulfonate salt (Tricaine, Cat#A5040; Sigma) in E3 embryo medium in a 50 μL glass capillary. Heart beat was stopped with 30 mM 2,3-Butanedione monoxime (BDM, Cat#B0753; Sigma) as indicated in individual experiments.

Dendra2 photoconversion experiments were performed on an inverted Zeiss LSM710 confocal microscope with the Plan-Apochromat 20x/0.8 M27 or LD LCI Plan-Apochromat 25x/0.8 Imm Korr DIC M27 objectives. Embryos were embedded in 1% low-melting point agarose and 0.016% Tricaine in E3 embryo medium in glass-bottom plates orienting the anterior dorsal side of the embryo towards the bottom of the plate.

Panoramic SPIM and high-resolution SPIM of the beating zebrafish heart as well as image processing (Mercator projections and reconstruction of the beating heart) were performed essentially as described (Mickoleit et al., 2014; Schmid et al., 2013).

Images were processed using Leica LAS, ImageJ/Fiji, Imaris, and Photoshop CS6. The area of the BA, ventricle and regions positive for *drl*:EGFP were measured from confocal image Z-projections of dissected hearts using ImageJ/Fiji. Extracted data were processed with GraphPad Prism 7 using unpaired t-test with Welch correction, reported P values are two-tailed.

### Chemical treatments

SU5402 (Sigma) was administered to embryos at concentrations ranging from 2-5 μM in E3 at desired stages and for specific periods of time as indicated in the text. The drug was washed out through several washing steps with E3. Diaminorhodamine-4M AM solution (DAR-4M, Cat#D9194; Sigma) was diluted 1:1000 in E3 containing 0.003% 1-phenyl-2-thiourea (PTU, Cat#P7629; Sigma) and live embryos incubated in the staining solution for at least 48h at 28°C. DAR-4M was washed out through several washing steps before imaging. Controls were treated with equivalent amounts of DMSO.

## Acknowledgements

We thank Sibylle Burger and Seraina Bötschi for technical and husbandry support, Dr. Stephan Neuhauss for zebrafish support, the ZBM at UZH for imaging support, and the members of the Mosimann lab for constructive input. This work has been supported by a Swiss National Science Foundation (SNSF) professorship [PP00P3_139093] and SNSF R’Equip grant 150838 (Lightsheet Fluorescence Microscopy), a Marie Curie Career Integration Grant from the European Commission [CIG PCIG14-GA-2013-631984], the Canton of Zürich, the UZH Foundation for Research in Science and the Humanities, and the Swiss Heart Foundation to C.M; the Helmholtz Young Investigator Program VH-NG-736, Deutsche Forschungsgemeinschaft (DFG) PA2619/1-1, and Marie Curie Career Integration Grant from the European Commission (WNT/CALCIUM IN HEART-322189) to D.P; a Company of Biologists and EuFishBioMed travel grant to K.D.P.

## Author contributions

A.F., K.D.P., A.M., D.P., and C.M. designed, performed, and analyzed the experiments; K.D.P. performed panoramic SPIM under guidance of J.H.; M.M. performed and analyzed experiments in Fig. 3A-E and associated videos as supervised by J.H.; E.C.B. performed lineage trace experiments in Supplementary Figure 2; D.P. and C.M. supervised the project; A.F., D.P., and C.M. compiled data and wrote the manuscript.

## Competing financial interests

The authors declare no competing financial interests.

## Supplementary Material

**Supplementary Table 1: Comparison of *tbx1*:EGFP-transgenic zebrafish lines.**

Comparison of fluorescence intensity, the number of transgenic insertions and unspecific signals in F1 *tbx1:EGFP line I-VI* transgenics. Lines *I* (*Tg(-3.2tbx1:EGFP_I)^zh702^), IV*, and *V* were retained and all experiments performed in F2 generations or beyond of line *I*, and some experimental outcomes confirmed in the other retained lines.

**Supplementary Figure 1: *tbx1* reporter- and endogenous expression in different transgenic lines and at different timepoints.**

(**A,B**) Anterior expression in different transgenic *tbx1:EGFP* lines at 20 ss (**A**) and 24-28 hpf (**B**); lateral views, anterior to the left. Expression in all lines is confined to the anterior CPF with some minor unspecific signals in the notochord or scattered posterior skeletal muscles (see Table S1). (**C**) Cardiac *tbx1*:EGFP expression in all lines is strongest in the ventricle and outflow tract at 60 hpf; lateral view of the right side of the embryo due to better visibility of the outflow tract, anterior to the right. (**D,E**) Endogenous *tbx1* expression confines to a dorsal domain during gastrulation; (**D**) animal view, (**E**) lateral view; V: ventral, D: dorsal, A: animal, Veg: vegetal. (**F-H**) By mRNA ISH, *tbx1:EGFP* and *tbx1* expression is detected in anterior pharyngeal precursors from mid-somitogenesis to heart tube stages (16 ss to 30 hpf). Additionally, *tbx1* reporter expression marks the migrating heart field at the midline (asterisk) and emerging fin bud precursors (arrowheads); dorsal views, anterior to the top. (**I**) *tbx1*:EGFP and *tbx1* are confined to anterior domains with no expression in posterior structures; asterisks heart tube in *tbx1*:EGFP, arrowhead fin bud precursors; lateral views, anterior to the left. Scale bars 100 μm (**F-H**), 250 μm (**A,C**), 500 μm (**B,I**).

**Supplementary Figure 2: Transverse sections of *tbx1* reporter expression and CreERT2/*lox*-mediated *tbx1* lineage tracing.**

(**A-L**) Transverse agarose sections of 3 dpf transgenic or control embryos aligned top to bottom from anterior to posterior with endogenous EGFP reporter fluorescence (green) and nuclear stain (DAPI, blue); top-down confocal Z-sections; dorsal to the top. (**A**) In the anterior-most sections, *tbx1:EGFP* reporter expression is confined to neural crest-derived mandibular cartilage (m) and ALPM-derived cranial vasculature (cv). (**B,C**) Further posterior branchial arches (ba, containing mesodermal, endodermal and neural crest lineages) and the endodermal pharynx (ph) express *tbx1*:EGFP. (**D,E**) *tbx1*:EGFP reporter expression can be also found in the endodermally derived swim bladder (sb) and in the anterior gut (ag), and weakly in the liver (l) but not in the posterior gut segments (pg) at the height of the swim bladder, suggesting absence of *tbx1* reporter activity in more posterior segments. (**F**) No organ-specific EGFP signal can be observed in non-transgenic (wildtype, *wt*) controls, but autofluorescent signal in the skin and blood can be observed in controls and throughout all sections (asterisk **A-L**). (**G-K**) Akin to *tbx1*:EGFP reporter expression, *tbx1:creERT2*-mediated lineage tracing throughout gastrulation (shield-bud, tbx1>EGFP) marks the branchial arches (ba), pharynx (ph), anterior gut (ag), liver (l), and swim bladder (sb). As observed by live imaging of lineage-traced embryos (Figure 2), ventricular myocardium of the heart (h) is also marked. The posterior gut was only labelled in 2/7 embryos, supporting loss of enhancer activity in more posterior endodermal precursors. (**L**) No unspecific switching activity was observed in heat shock (hs) controls. Scale bars 500 μm.

**Supplementary Figure 3: Tissue-specific *tbx1* reporter expression and lineage tracing in cardiopharyngeal tissues.**

(**A**) Maximum intensity projections of fixed whole-mount immunofluorescence from 60 hpf *tbx1*:EGFP embryos counterstained with anti-EGFP and anti-MHC; ventral view, anterior to the top. *tbx1*:EGFP-positive cranio-branchial derivatives observed through lineage tracing and reporter expression in previous experiments include the cranial muscles inferior oblique (io) and adductor mandibulae (am). (**B**) *tbx1:creERT2* transgenics were crossed to *ubi:Switch* and induced with 4-OHT at the end of gastrulation (90% epiboly). *ubi:Switch* (*ubi:loxP-EGFP-loxP-mCherry*) switches from EGFP to mCherry expression in all cells expressing the *creERT2* transgene at the time of 4-OHT induction. (**C**) mCherry-expressing *tbx1:creERT2* derivatives (*tbx1*>mCherry) at 4.5 dpf can be found in the heart (h), cranio-branchial derivatives (cbd), mandibular cartilage (m), and in the ear (ea). The inlet in (**C’’**) depicts the brightfield image of (**C-C’’**); lateral view, anterior to the left. (**D**) *tbx1:creERT2* or *drl:creERT2* transgenics, respectively, were crossed to the ubiquitous *hsp70l:Switch loxP* tracer line, 4-OHT-induced at shied stage or 90% epiboly, and heat-shocked at 3 dpf. (**E,F**) Maximum intensity projections from SPIM-imaged 3.5 dpf embryo; ventral views, anterior to the top. (**D**) *tbx1:creERT2;hsp70l:Switch* double-transgenic embryos were induced with 4-OHT at shield stage to capture the earliest derivatives of *tbx1* reporter-expressing progenitors. *tbx1* lineage-traced cells (*tbx1*>EGFP) are located in ventricular (V) myocardium, atrial (A) myocardium at the inflow tract, and in the DAR-4M-marked bulbus arteriosus (BA). Additionally, cranio-branchial derivatives (cbd) and cranial vessels (cv) are readily marked by *tbx1* lineage tracing. (**B**) *drl:creERT2*-mediated lineage tracing (*drl*>EGFP) was initiated at 90% epiboly to label LPM derivatives that can be found in all cardiac structures (V, A, BA) and cranial vasculature (cv). Scale bars 100 μm (**A**,**E,F**), 250 μm (**C**).

**Supplementary Figure 4: *tbx1* reporter-expression at the base of the linear heart tube and smooth muscle of the BA.**

(**A-C**) Maximum intensity projections of representative stages from SPIM-imaged *tbx1:EGFP;myl7:DsRed2* double transgenic embryos, dorsal views, anterior to the top. Imaging was initiated at 18 hpf and heart development was followed up to 30 hpf. *tbx1*:EGFP-only cells were observed at the base of the *myl7*:DsRed2-positive linear heart tube at all stages imaged (arrow heads). Imaging was performed on *tbx1:EGFP line IV* embryos; note the strong notochord expression (asterisk) not observed in *line I* (see also Figure S1 and Table S1). (**D,E**) Transverse agarose sections of *tbx1:EGFP;drl:mCherry* (**D**) or *drl:mCherry* (**E**) transgenic embryos with endogenous EGFP (green) and mCherry (magenta) reporter fluorescence and nuclear stain (DAPI, blue); top-down confocal z-sections; dorsal to the top. *drl:mCherry* is expressed in endothelial cells (ec) and blood (b), *tbx1*:EGFP is specific to the smooth muscle layer of the bulbus arteriosus (sm). *tbx1* reporter expression can be also detected in the myocardium of the inner curvature (cm) and in the branchial arches (ba). Scale bars 50 μm.

**Supplementary Figure 5: Myocardium in all ventricular segments derives from the *tbx1*:Dendra2-expressing sheath at the base of the forming heart tube.**

(**A-C**) Maximum intensity projections of the heart shown in Figure 5B with myl7:AmCyan demonstrating that photoconverted cells from the *tbx1* reporter-expressing sheath at the base of the forming heart tube at 22 hpf contribute to ventricular myocardium at 3.5 dpf; ventral view, anterior to the top. (**D**) A second representative tbx1:Dendra2 embryo converted at the base of the forming heart tube at 22 hpf is shown; maximum intensity projection, dorsal view, anterior to the top. (**E-H**) Live SPIM-imaging of the same embryo depicted in (**D**) at 3.5 dpf; ventral view, anterior to the top. Optical z-sections from ventral (**E**), medial (**F**) and dorsal (**G**) regions as well as the maximum intensity projection (**H**) show that regions of the ventricular myocardium, including most distal segments, but not the bulbus arteriosus, are contributed from the photoconverted sheath at 22 hpf. Dendra2-red in all or only the distal-most ventricular myocardium (as depicted in **A-C**) was observed in 10/11 embryos with *tbx1* reporter-expressing sheath conversion at 22 hpf. In 1/11 embryos, no Dendra2-red cells could be found in the entire anterior region of the embryo. Scale bars 50 μm.

**Supplementary Figure 6: Different regions of the pharyngeal LPM contribute to distinct parts of the BA and craniofacial structures.**

(**A-D**) Maximum intensity projections and schematics of representative photoconverted tbx1:Dendra2 embryos; dorsal views, anterior to the top. At 22 hpf, *tbx1*:Dendra2-expressing embryos were illuminated with UV light in a confined region of interest to convert Dendra2-green to Dendra2-red in specific *tbx1* reporter-expressing domains. (**E-H**) Maximum intensity projections and graphical representations of hearts of tbx1:Dendra2 embryos photoconverted in (**A-D**) at 3.5 dpf with marked contribution to the heart by photoconverted cells; ventral views, anterior to the top. (**A,E**) Photoconversion of a broad area in the *tbx1*:Dendra2-positive pharyngeal LPM posterior to and on the left of the cardiac cone marks the left side of the BA (dotted outline). (**B,F**) The most medial region of the area converted in (**A**) contributes to the most proximal part of the BA suggesting that these progenitors migrate and add the earliest to the heart. (**C,G**) A lateral and anterior region to (**B**) contributes to the medial part of the BA. (**D,H**) The most posterior and lateral region makes the distal part of the BA and contributes to cells of the ascending aorta. **(I-K)** Maximum intensity projections of photoconverted tbx1:Dendra2 embryos at 3.5 dpf; ventral views, anterior to the top. Photoconverted regions as in (**B-D**), (**J**) and (**K**) depict the same embryos as shown in (**C**) and (**D**), respectively; (**I**) shows an embryo that was photoconverted in an equivalent but right-hand sided region as shown in (**B**). Progenitors of different parts of the BA derive from the same areas as different parts of craniofacial muscles and cartilage. (**L**) Maximum intensity projections of an unconverted *tbx1*:Dendra2 embryo; ventral view, anterior to the top. Red signal from blood (asterisks) and on top of the pericardium (arrows) were observed in unconverted Dendra2 embryos imaged at 3.5 dpf using lightsheet microscopy, suggesting unspecific auto-fluorescence in these structures picked up at the imaging conditions used for Dendra2 experiments. Scale bars 50 μm.

**Supplementary Figure 7: Aberrant formation of the DAR-4M-stained BA upon FGF signaling perturbation.**

**(A-D)** *myl7*:EGFP, DAR-4M-stained embryos treated with DMSO or 5μM SU5402 during (14 ss-22hpf) or after (24 hpf-34 hpf) heart tube formation; lateral views, anterior to the left. Absent BA formation can only be observed in embryos treated with SU5402 from mid-somitogenesis to heart tube stages (arrow head **B**), but not when signaling inhibition is initiated at 24 hpf (arrow head **D**). Scale bars 100 μm.

**Supplementary Video 1: Panoramic lightsheet-imaging of the early development of a *tbx1*:EGFP;dr/:mCherry embryo.**

Mercator projection of early developmental stages from 8 -15 hpf, *tbx1*:EGFP is depicted in green, *drl*:mCherry in magenta, dorsal view, anterior to the left. Movie stacks were acquired every 2 min. Frame rate of the movie: 29 frames per second (fps). 1 sec in the movie corresponds approximately to 30 min in development.

**Supplementary Video 2: Lightsheet imaging of the cardiopharyngeal field of a *tbx1*:EGFP;*drl*:mCherry embryo.**

Maximum intensity projection with the dorsal view of the anterior region of a *tbx1*:EGFP;*drl*:mCherry transgenic, anterior to the top right corner; *tbx1*:EGFP depicted in green, *drl*:mCherry in magenta. Cardiac development was followed from 14 ss to 23 hpf. Movie stacks were acquired approximately every 7.5 min. Frame rates 7 fps. 1 sec in the movie corresponds approximately to 52.5 min in development.

**Supplementary Video 3: Animated 3D segmentation of the *tbx1* reporter-expressing sheath at the base of the heart tube.** Rotation of the 3D segmentation of still from Video S2 shown in Figure 3D revealing a *tbx1* reporter-expressing sheath of cells at the base of the forming heart tube and engulfing the *drl* reporter-expressing endocardium at 22-23 hpf.

**Supplementary Video 4: Lightsheet-imaging of the developing heart tube of a *tbx1*:EGFP;*myl7*:DsRed2 embryo.**

Maximum intensity projection with the dorsal view of the anterior region of a *tbx1*:EGFP;*myl7*:DsRed2 transgenic, anterior to the top; *tbx1*:EGFP depicted in green, *myl7*:DsRed2 in magenta. Heart tube development was followed from 18 ss to 30 hpf. Movie stacks were acquired approximately every 2.8 min. Frame rates 10 fps. 1 sec in the movie corresponds approximately to 28 min in development.

**Supplementary Video 5: Time-lapse of a high-speed SPIM-imaged beating heart in a *tbx1*:EGFP;*myl7*:DsRed2 transgenic.**

Heart development was followed from 28-52 hpf, all time points are synchronized and shown at contraction phase 27, heart imaged from right side of the embryo, lateral view, anterior to the top, ventricle to the upper left, atrium to the lower right. *tbx1*:EGFP is depicted in cyan, *myl7*:DsRed2 in red. Frame rates 7 fps. 1 sec in the movie corresponds approximately to 144 min in development.

**Supplementary Video 6: *tbx1*:EGFP-expression in a beating heart at 28 hpf reconstructed from highspeed SPIM-imaging.**

Volume rendering showing a cardiac cycle in slow motion in a 28 hpf embryo heart as imaged in Video S5. Only EGFP expression is shown to highlight that all *tbx1* reporter-expressing cardiac cells move during cardiac contraction.

